# Pyrimidine salvage in *Toxoplasma gondii* as a target for new treatment

**DOI:** 10.1101/2023.11.01.565095

**Authors:** Hamza A. A. Elati, Amber L. Goerner, Bruno Martorelli Di Genova, Lilach Sheiner, Harry P. de Koning

## Abstract

Toxoplasmosis is a common protozoan infection that can have severe outcomes in the immunocompromised and during pregnancy, but treatment options are limited. Recently, nucleotide metabolism has received much attention as a target for new antiprotozoal agents and here we focus on pyrimidine salvage by *Toxoplasma gondii* as a drug target. Whereas uptake of [^3^H]-cytidine and particularly [^3^H]-thymidine was at most marginal, [^3^H]-uracil and [^3^H]-uridine were readily taken up. Kinetic analysis of uridine uptake was consistent with a single transporter with a K_m_ of 3.3 ± 0.8 µM, which was inhibited by uracil with high affinity (K_i_ = 1.15 ± 0.07 µM) but not by thymidine or 5-methyluridine, showing that the 5-Me group is incompatible with uptake by *T. gondii*. Conversely, [^3^H]-uracil transport displayed a K_m_ of 2.05 ± 0.40 µM, not significantly different from the uracil K_i_ on uridine transport, and was inhibited by uridine with a K_i_ 2.44 ± 0.59 µM, also not significantly different from the experimental uridine K_m_. The reciprocal, complete inhibition, displaying Hill slopes of approximately ∼1, strongly suggest that uridine and uracil share a single transporter with similarly high affinity for both, and we designate it uridine/uracil transporter 1 (TgUUT1). While TgUUT1 excludes 5-methyl substitutions, the smaller 5F substitution was tolerated as 5F-uracil inhibited uptake of [^3^H]-uracil with a K_i_ of 6.80 ± 2.12 µM (*P* > 0.05 compared to uracil K_m_). Indeed, we found that 5F-Uridine, 5F-uracil and 5F,2’-deoxyuridine were all potent antimetabolites against *T. gondii* with EC_50_ values well below that of the current first line treatment, sulfadiazine. *In vivo* evaluation also showed that 5F-uracil and 5F,2’-deoxyuridine were similarly effective as sulfadiazine against acute toxoplasmosis. Our preliminary conclusion is that TgUUT1 mediates potential new anti-toxoplasmosis drugs with activity superior to the current treatment.

## 1. Introduction

*T. gondii* has the ability to infect any nucleated cell of any warm-blooded animals or bird, including those of humans, giving it possibly the widest host range of any parasite (Tenter et al., 2000). *T. gondii*, being acquired orally and not requiring a vector, is not geographically limited and is estimated to have infected approximately one-third of the global human population (Hill et al., 2005, Montoya and Liesenfeld, 2004). It can remain latent in the majority of its hosts, but can be deadly in immunocompromised people, and create severe adverse impacts on the unborn if acquired during pregnancy. Acute toxoplasmosis can result in severe cerebral outcomes including retinochoroiditis, encephalitis, hydrocephalus, convulsions and intracerebral calcification (Hill et al., 2005). In most immunocompetent humans, however, toxoplasmosis does not need treatment and protective immunity is naturally acquired.

However, treatment is vital for toxoplasmosis patients that are pregnant or immunocompromised. There are very few drugs for toxoplasmosis treatment which, moreover, have serious adverse effects and they are not useful for the bradyzoite stage. The first-line treatment for acute toxoplasmosis is the combination of two antimicrobial antifolate agents: pyrimethamine (PYR) or trimethoprim, which inhibit dihydrofolate reductase (DHFR) and sulphonamides such as sulfadiazine, sulfamethoxazole or sulfadoxine, which inhibit dihydropteroate synthetase (DHPS). The combination acts synergistically to disrupt the parasite’s folate metabolism (Alday and Doggett, 2017; Dunay et al., 2018). However, as pyrimethamine also inhibits the human DHFR (Heppler et al., 2022) it can cause haematological toxicity, making it necessary to give leucovorin or folinic acid together with the first-line treatment to combat the side effect (Alday and Doggett, 2017; Dunay et al., 2018).

Folate is an essential cofactor for the biosynthesis of thymidine nucleotides, and nucleotide metabolism in protozoan parasites is a vital biological process and increasingly explored as a drug target. Nucleoside antimetabolites have been shown to be effective drug candidates against various protozoan parasites, such as *Trichomonas vaginalis* (Natto et al., 2021a, Natto et al., 2021b), *Trypanosoma brucei* (Hulpia et al., 2020a, Hulpia et al., 2020b), *Trypanosoma congolense* (Mabille et al., 2022), *Trypanosoma vivax* (Ungogo et al., 2023), *Trypanosoma cruzi* (Fiuza et al., 2022) and apicomplexan parasites such as *T. gondii* (Campagnaro et al., 2022; Al Safarjalani et al., 2010; Kim et al., 2010) and *Plasmodium falciparum* (Cheviet et al., 2019).

Most protozoan parasites including *T. gondii* are able to synthesise pyrimidine nucleotides *de novo* as well as salvage pyrimidines such as thymidine and uracil (Ali et al., 2013; De Koning et al., 2003; Ghosh and Mukherjee, 2000; De Koning and Jarvis, 1998; Gudin et al., 2006; Finley et al., 1988; Papageorgiou et al., 2005) but some exceptions exist. *Giardia lamblia*, *Tritrichomonas foetus*, *T. vaginalis* (Hassan and Coombs, 1988; Berens et al., 1995) and *C. parvum* (Rider and Zhu, 2010) are pyrimidine auxotrophs and rely on pyrimidine uptake from the environment. In contrast, *P. falciparum* is incapable of pyrimidine salvage (Gutteridge et al., 1970) but fully competent to synthesise pyrimidine nucleosides *de novo* (Polet and Barr, 1968).

UMP is the final product of pyrimidine biosynthesis, from which all other pyrimidine nucleotides, nucleosides and nucleobases can be produced. In addition, *T. gondii* is able to acquire uracil, (2’-deoxy)uridine and (2’-deoxy)cytidine from the host cell and transform the nucleosides to uracil using cytidine deaminase and uridine phosphorylase, and subsequently to UMP, exclusively by uridine phosphoribosyltransferase (UPRT) activity (Iltzsch, 1993; Fox and Bzik, 2020). This pyrimidine salvage strategy is known as a “salvage funnel”, routing all salvaged pyrimidines to uracil and thence to UMP (Pfefferkorn, 1988).

Purine and pyrimidine transporters have been extensively studied in kinetoplastids, particularly in *Leishmania* and *Trypanosoma* species (Aldfer et al., 2022b; Aldfer et al., 2022a; Ungogo et al., 2023; Al-Salabi et al., 2003) as well as the apicomplexan *P. falciparum* (Quashie et al., 2008; Deiskin et al.; 2015; Campagnaro and De Koning, 2020; Parker et al., 2000) but nucleoside and nucleobase transport in *T. gondii* has received relatively scant attention. However, a few studies have been conducted on purine uptake by extracellular *T. gondii* tachyzoites and these found a low affinity high-capacity adenosine transporter, which was named TgAT1 (Schwab et al., 1995), and was later supplemented with a high-affinity adenosine/inosine transport activity, TgAT2, and a separate high-affinity purine nucleobase (hypoxanthine/guanine) transporter, TgNBT1 (De Koning et al., 2003). The gene encoding TgAT1, Tg_244440, was identified from an adenine arabinoside (Ara-A)-resistant cell line, was characterized by expression in *X. laevis* oocytes, and was confirmed to be a low affinity, high-capacity transporter of adenosine (K_m_ of 114 µM) (Chiang et al., 1999). Recently, expression of Tg_244440/TgAT1 in a *T. brucei* cell line provided further confirmation of the low affinity for adenosine but additionally found that the carrier displays higher affinity for oxopurine nucleobases and nucleosides (Campagnaro et al., 2022).

However, the transport of pyrimidines by *T. gondii* has not yet been reported and so far the only information is that uridine and thymidine appear to be inhibitors, and thus perhaps substrates, of TgAT2 (De Koning et al., 2003). Yet it is clear that at least some pyrimidines can be salvaged by intracellular trophozoites and incorporated into nucleic acids (Pfefferkorn and Pfefferkorn, 1977). Indeed, the intracellular growth of pyrimidine auxotroph lineages can be rescued with a high concentration of uracil and uridine in the culture medium, providing conclusive proof that at least those pyrimidines can be salvaged by intracellular trophozoites (Fox and Bzik, 2002; Fox and Bzik, 2010). In contrast, thymidine is not salvageable by *T. gondii*, as it lacks thymidine kinase activity (Fox et al., 2001). Here, we investigate the uptake of pyrimidine nucleosides and uracil in isolated *T. gondii* tachyzoites and identify a previously unreported uridine/uracil carrier in *T. gondii* tachyzoites that appears to facilitate the uptake of potent pyrimidine antimetabolites.

## 2. Methods

### 2.1 Host cells and parasites

The F3 (RH Δku80 TATi) strain (Sheiner et al., 2011) was first made fluorescent red through random integration of the tubTandemTomatoRFP/sagCAT (pCRT2t) plasmid (Chtanova et al., 2008) and selection via flow cytometry. This cell line was always used as the primary cell line for all transport assays and drug screening assays in this study.

### 2.2 *T. gondii* tachyzoites cell culture

*T. gondii* tachyzoites were cultured in human foreskin fibroblasts (HFF), sourced from ATCC (SCRC-1041). HFFs and parasites were culture in Dulbecco’s Modified Eagle’s Medium (DMEM), containing 4.5 gL^-1^ glucose, supplemented with 10% (v/v) fetal bovine serum, 4mM L-glutamine and penicillin/streptomycin antibiotics and grown at 37°C with 5% CO_2_. When needed anhydrotetracycline (ATc) was added to the medium at a final concertation of 0.5 µM.

### 2.3 Radiolabelled and chemical compounds

The following tritium radiolabelled compounds were used during the project: [^3^H]-Uracil (40 Ci/mmol), [^3^H]-Thymidine (71.7 Ci/mmol), [^3^H]-Uridine (60 Ci/mmol) were obtained from PerkinElmer (Waltham, MA, USA). [^3^H]-Cytidine (20 Ci/mmol) was purchased from American Radiolabelled Chemicals Incorporated (St Louis, MO, USA). Uridine, Uracil, Thymidine, 5-Floururacil, 5-Flourouridine, 2’-Deoxyuridine, 3’-deoxythymidine, Sulfadiazine, NBMPR, Adenosine, Inosine, resazurin sodium salt and Phenyl Arsine Oxide were bought from Sigma-Aldrich (Poole, UK). 5-Flouro 2’Deoxyuridine was from VWR; 5-Methyluridine was from Alfa Aesar; 2’,3’-dideoxyuridine was from Carbosynth; Pyrimethamine was from Fluka; Ara-A was from ICN Biomedicals.

### 2.4 Drug sensitivity assay for *T. gondii* tachyzoites

The *in vitro* inhibitory effects of antimicrobial nucleoside agents on *T. gondii* growth were determined in HFF cell cultures using 96-well black plates (Optical-Bottom Plates with Polymer Base, ThermoFisher Scientific). Sulfadiazine (positive control) and the nucleoside compounds were already dissolved in DMSO at known concentrations, usually prepared as 10 mM. After that, every compound was diluted in DMEMc at 4x final concentration. 100 µL of 4x compounds were transferred to the wells of the first column of the 96-well black plate in triplicate or duplicate. The compounds were mixed, and 100 µL was transferred using the multichannel pipette to the next column before being transferred to each column through the entire plate over one row, leaving the last column as a negative control without drug. After that, naturally, freshly egressed parasites were collected and filtered through a 3-µm filter (Sigma-Aldrich). ∼1000 parasites/well were added and incubated for 6 days at a 37 °C/5% CO_2_ in a humidified incubator. On day 6, a PHERAstar plate reader (BMG Labtech, Germany) was used to read the fluorescence intensity at 540 nm for excitation and 590 nm for emission. The EC_50_ values and fluorescence data were determined and plotted by using Prism 8.0 (GraphPad). Each experiment was independently performed 3-5 times.

### 2.5 Drug cytotoxicity assay for HFF cells using Alamar blue dye

HFF cytotoxicity assay was done exactly as described above in **the drug sensitivity assay for *T. gondii* tachyzoites**, except that no parasites were added during this assay. Phenyl Arsine Oxide (PAO) was always used as a positive control. The plates were incubated at 37 °C under 5% CO_2_ in a humidified incubator for 6 days to be consistent with the drug screening assay against *T. gondii*. On day 6, 10 µL of a 12.5 mg/100 mL ddH_2_O resazurin solution was added to each well. In addition, 200 µL of DMEMc media was placed in one row in a new 96-well plate without any host cells to determine the level of background fluorescence for the calculation, and 10 µL of a 12.5 mg/100 mL ddH_2_O resazurin solution was added to those wells as well. The plates were incubated for 3-4 h at 37 °C and 5% CO_2_ in a humidified incubator. Subsequently, fluorescence was read using the PHERAstar plate, at λ_exc_ 540 nm and λ_em_ 590 nm. The data was plotted using an equation for a sigmoid curve with variable slope (4 parameter) using Prism 8.0 (GraphPad); EC50 values were extrapolated if >50% was achieved at the highest test compound concentration. Each biological experiment was performed on 3-5 independent occasions. The background plate result was subtracted from the reading from the raw data result of the fluorescent intensity of the test compounds.

### 2.6 Transport studies using extracellular *T. gondii* tachyzoites

Transport of pyrimidine nucleosides/bases (uridine, uracil, thymidine and cytidine) into extracellular *T. gondii* tachyzoites was performed using a modified oil-stop technique previously described for transport measurements with *T. gondii* tachyzoites (De Koning et al., 2003), with only minor modification and optimisation in the oil mixture. *T. gondii* tachyzoites were cultured in a confluent HFF T175 vented flask kept at 37 °C and 5% CO_2_ in a humidified incubator until the tachyzoites parasites became extracellular and the extracellular tachyzoites culture suspension was collected and filtered through 47 mm diameter, 3 µM pore-size Nuclepore polycarbonate filters (Whatman International Ltd, UK) to remove the HFF debris, and centrifuged at 1500 x *g* for 15 min at 4 °C. The assay buffer (AB; 33 mM HEPES, 98 mM NaCl, 4.6 mM KCl, 0.5 mM CaCl_2_, 0.07 mM MgSO_4_, 5.8 mM NaH_2_PO_4_, 0.03 mM MgCl_2_, 23 mM NaHCO_3_, 14 mM D-glucose, pH 7.3) was used to wash the pellets twice. Tachyzoites were counted using a Neubauer haemocytometer (Weber Scientific Ltd, UK) and resuspended in AB at a density of 2 × 10^8^ cells/mL; the parasites suspension was allowed to recover from centrifugation stress for 30 min at room temperature (RT). Next, 100 µL of the AB containing radiolabelled nucleoside/nucleobase test compound at a known concentration, was layered over 200 µl of an oil mixture (1:5 mixture of mineral oil (Sigma) and di-*n*-butyl phthalate (BDH Ltd, UK) for tachyzoites cultures in a microfuge tube. Centrifugation was briefly applied to allow the aqueous layer with radiolabel and test compound to be layered perfectly on top of the oil. Transport assays, performed at RT, were started by adding 100 µL of the tachyzoite cell suspension in AB to the mixture of oil and radiolabel test compound, with a predetermined incubation time, taking care that the aqueous layers mixed and remained on top of the oil. Upon completion of incubation time, 750 µL of ice-cold stop solution was added to stop the transport assay or to saturate the transport. An ice-cold solution of unlabelled substrate was used as a stop solution using a concentration between 0.5 mM to 2 mM, depending on the saturation concentration of the transporter and the aqueous solubility of the compound. This step is immediately followed by centrifugation at 14800 × *g* for 1 min to pellet the cells under the oil layer. The microfuge tube was flash-frozen in liquid nitrogen, and the bottom of each tube, containing the cell pellet, was cut off and collected in 6 mL scintillation vials (LabLogic Systems Ltd, Sheffield, UK). The pellets were lysed using 300 µL of 2% (w/v) sodium dodecyl sulfate (SDS; 2% SDS in H_2_O) at RT for at least 40 min on a rocking platform (GENEO Biotech Products GmbH, Germany). To each vial a volume of 3 mL scintillation fluid (Optiphase Hisafe-2) was added, and the vials were left overnight in a dark area at RT. Vigorous shaking was done before the scintillation vials were measured using a Hidex 300 SL liquid scintillation counter (LabLogic Systems Ltd, Sheffield, UK). The assays were performed three times in independent experiments, each in triplicate.

### 2.7 Pyrimidine drug mouse assay

24 C57BL/6J mice (The Jackson Laboratory) were infected intraperitoneally (i.p.) with 1000 Me.49.b7 *T. gondii* parasites diluted in 200 µl of 1×PBS. Body weights were determined prior to infection and once a day until euthanasia at day 10. Starting 72 h post infection, groups of 6 mice received daily drug treatment, spaced exactly 24 h, through day 9, or served as untreated control: 5-fluoro-2’deoxyuridine (5-F,2’dUrd; 25 mg/kg i.p., 200 µl in water), 5-fluorouracil (5-FU, 25 mg/kg i.p., 200 µl in water); Sulfadiazine (100 μg/L, oral in water as described (Saeji et al., 2005)); control mice were injected with 200 µl water i.p.. Following euthanasia by CO_2_ and cervical dislocation as per IUCUC guidelines, the lungs were dissected, removed and stored at −80 °C until DNA extraction.

Daily animal health monitoring began before infection and lasted through the experiment. Mice were euthanized if predetermined welfare thresholds were exceeded. All animal experiments were carried out under University of Vermont Institutional Animal Care and Use Committee protocol PROTO202100038.

DNA was extracted using the Qiagen DNeasy kit using the manufacturer’s instructions and collected in 50 µl of elution buffer. DNA concentration was determined using a Nanodrop spectrophotometer (Thermo Fisher Scientific). Parasite burden in each lung tissue sample was determined using qPCR. 600 ng of each sample was used to normalize the relative burden within each organ and compared to a standard curve of isolated *T. gondii* parasites. SYBR green PCR Mastermix (Applied Biosystems) was used in a 20 µl volume, along with 500 nM of forward (5’-TCCCCTCTGCTGGCGAAAAGT-3’) and reverse (5’-AGCGTTCGTGGTCAACTATCG-3’) primer to the *T. gondii* B1 gene, as described (Vizcarra et al., 2023). The amplification program, ran on a Quant studio 5 thermocycler, was 50 °C for 2 min, 95 °C for 10 min then 40 cycles of 95 °C 10 s, 60 °C 30 s, followed by 60 °C for 30 s and 95 °C for 5 s.

Experiments were analysed and reviewed using the Design and Analysis 2nd edition software (Thermo Fisher). Normalization and final calculations used Microsoft Excel. The standard curve was generated by serial dilution of Me49 parasite genomic DNA. Determination of parasite burden per sample was calculated as pg of parasite genomic DNA, based on the standard curve, per 600 ng of host lung tissue.

## 3. Results

### 3.1 Thymidine and cytidine are not significantly salvaged by *T. gondii*, but uridine and uracil are

Sub-micromolar concentrations of radiolabelled pyrimidines were incubated over preset intervals with *T. gondii* tachyzoites and uptake was monitored. Uptake of 0.25 µM [^3^H]-thymidine over 10 min was barely detectable, although a line with a non-zero slope (*P* = 0.012, F-test) and non-significant deviation from linearity (*P* > 0.999, runs test) could be plotted (Figure 1A). However, this does not represent any mediated, high affinity uptake as the addition of 1 mM unlabelled thymidine did not significantly inhibit uptake over 10 min (*P* < 0.05, t-test, n=3). Likewise, 250 µM uridine did not inhibit the thymidine uptake and we conclude that if this does represent transporter-mediated thymidine uptake it must be very low affinity and not relevant at physiological concentrations. More likely, this represents non-specific accumulation of trace amounts of thymidine by a non-specific mechanism such as endocytosis.

**Figure 1.**
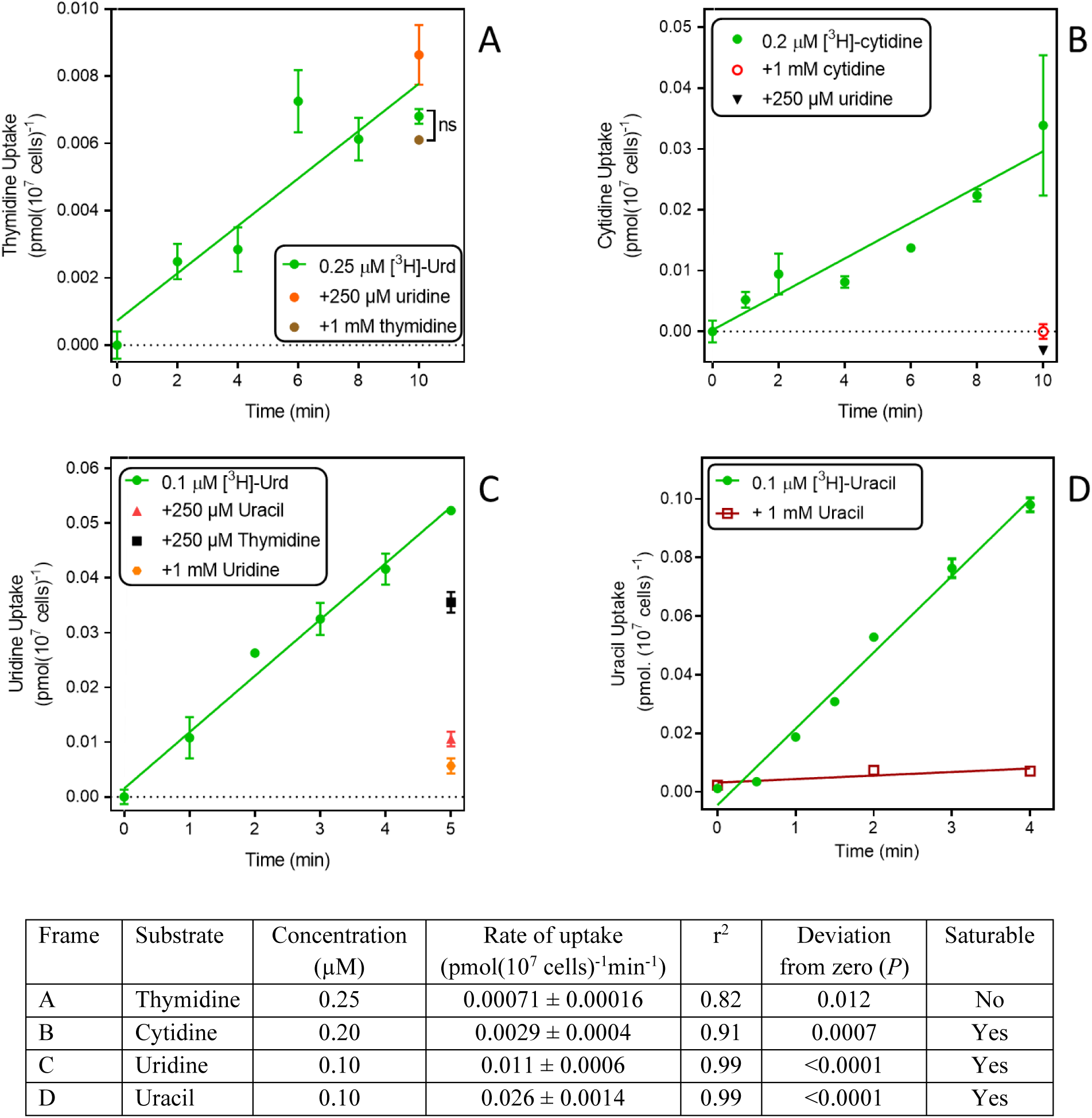
Uptake of pyrimidines by *T. gondii* tachyzoites. Uptake of tritiated thymidine (A), cytidine (B), uridine (C) and uracil (D) was determined over the indicated time intervals. Details of the graphs is shown in the accompanying table. All experiments were performed in triplicate, error bars are SE and, when not shown, fall inside the symbol. Lines were calculated by linear regression in Prism 8.0; none of these lines was significantly non-linear by F-test (P >0.05). For each frame, the analysis of the linear regression is presented in the table.

Uptake of 0.2 µM [^3^H]-cytidine was > 4-fold faster than thymidine and while this still represents a slow rate (0.0029 pmol(10^7^ cells)^-1^min^-1^), it was fully inhibited by the addition of either 1 mM unlabelled cytidine or 1 mM uridine (Figure 1B), showing that cytidine was taken up by a saturable transport system.

Uptake of [^3^H]-uridine was quite robust and easily allowed quantification at 0.1 µM over 5 min, with a rate of 0.010 pmol(10^7^ cells)^-1^min^-1^ (Figure 1C). This uptake appeared to be somewhat inhibited by 250 µM thymidine (*P* > 0.05), 80% by 250 µM uracil and 90% by 1 mM unlabelled uridine (both *P* < 0.01); the level of inhibition by uracil was not significantly different from that by uridine (*P* > 0.05, unpaired t-test).

Uptake of 0.1 µM [^3^H]-uracil was even more robust, at a rate of ∼2.5-fold that of uridine (0.026 pmol(10^7^ cells)^-1^min^-1^), and it was fully saturable, with uptake in the presence of 1 mM unlabelled uracil being not significantly different from zero (*P* = 0.37, F-test) (Figure 1D). We conclude that *T. gondii* tachyzoites do not express any thymidine transporters but do express one or more carriers for uracil and/or uridine and appear to have a minor capacity to take up cytidine. Since the cytidine uptake was fully inhibited by uridine, our data would be consistent with a model of a single transporter with the following substrate preference: uracil > uridine > cytidine >> thymidine.

### 3.2 Uridine and uracil share the same transporter

Uridine and uracil were both taken up with high affinity by tachyzoites. 0.1 µM [^3^H]-uridine was dose-dependently inhibited by unlabelled uridine in a mono-phasic way (Figure 2A), with an average K_m_ of 3.34 ± 0.82 µM and V_max_ of 1.21 ± 0.24 pmol(10^7^ cells)^-1^min^-1^ (n=3). Uridine transport was also inhibited by uracil, with an average K_i_ of 1.15 ± 0.07 µM (n= 3; not significantly different from the uridine K_m_ by t-test), and by 5-fluorouracil (5-FU) with a K_i_ of 5.24 ± 0.72 µM (n=3; *P* < 0.01 from uracil). These data are consistent with a model of a single uridine/uracil transporter that perhaps slightly favours uracil over uridine but is essentially a dual uridine/uracil transporter. Consistent with this model we found very similar parameters when using [^3^H]-uracil as the substrate (all n=3): K_m_ 2.05 ± 0.40 µM, K_i_ (uridine) 2.4 ± 0.59 µM, K_i_ (5-FU) 6.80 ± 2.12 µM (Figure 2B). None of these values was significantly different from the equivalent values using [^3^H]-uridine, which confirms that the tachyzoites express a uridine/uracil transporter, which we designate TgUU1.

**Figure 2.**
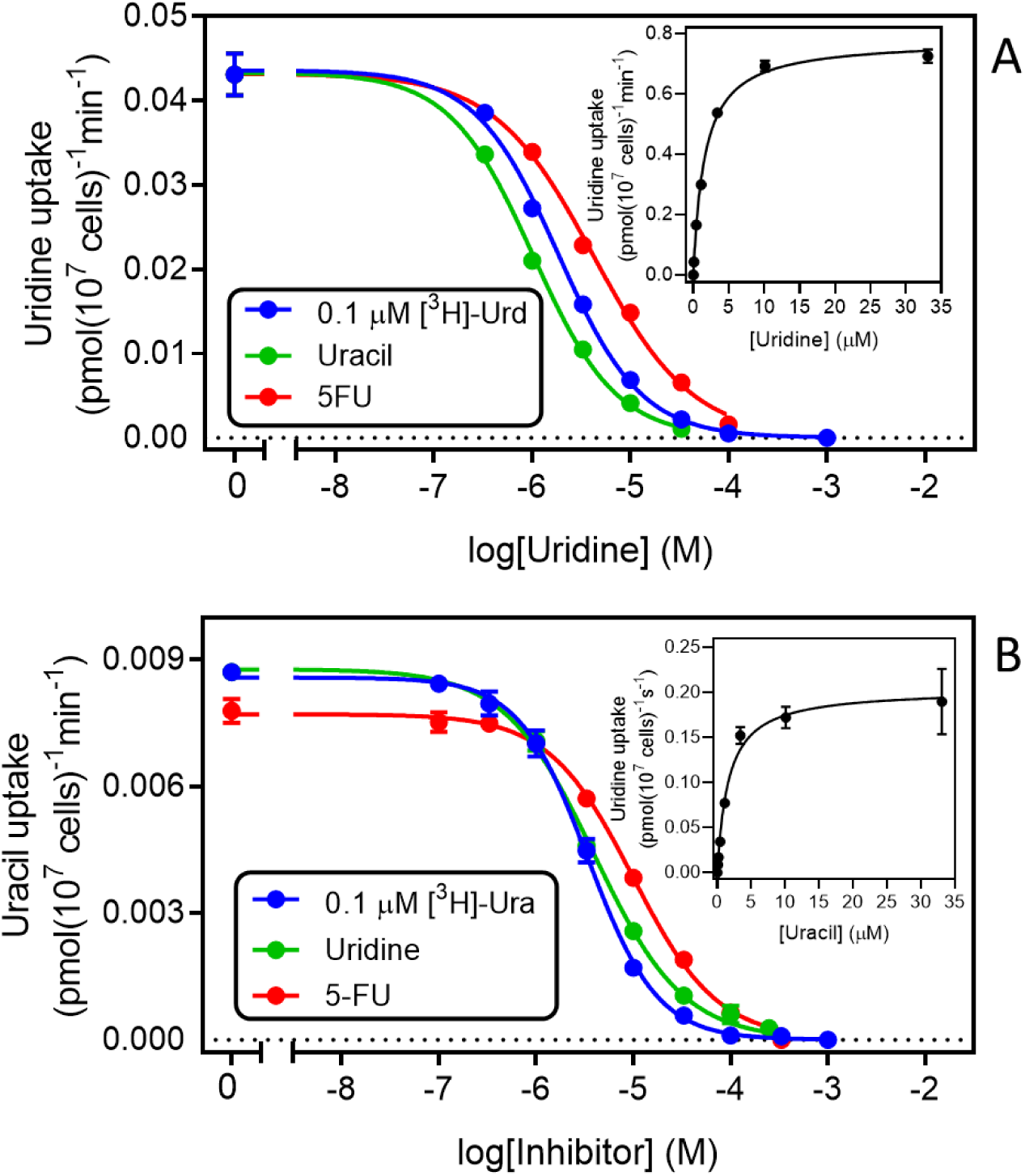
Kinetic parameters for the transport of uridine (A) and uracil (B) in *T. gondii* tachyzoites. Both frames depict a single representative experiment in triplicate, of dose-dependent inhibition of either 0.1 µM [^3^H]-uridine (frame A) or 0.1 µM [^3^H]-uracil. Error bars are SEM. Insets are the conversion of the inhibition of the inhibition of the uptake of radiolabelled permeant with unlabelled permeant to a Michaelis-Menten saturation plot. Hill slopes for the 6 plots varied between −1.2 and −0.86, consistent with a model for a single transporter.

Further characterisation of TgUU1 was conducted using 0.1 µM [^3^H]-uridine as the permeant. The transporter proved to be quite specific for pyrimidines, as inosine and adenosine, although able to inhibit uridine uptake, displayed much lower affinity than uracil, uridine and 5-FU (Figure 3A), with average K_i_ values of 28.2 ± 4.1 µM and 112 ± 6 µM, respectively (n=3). We next probed the reason for the high apparent selectivity for uridine over thymidine, and determined a K_i_ of 1010 ± 145 µM for thymidine, which corresponds to a change from −31.2 kJ/mol to −17.1 kJ/mol in Gibbs free energy of binding (δ(ΔG^0^) = 14.1 kJ/mol; Table 1). This is attributable to both differences between uridine and thymidine, being the presence of a 2’-OH group in uridine and 5-methyl in thymidine, as 2’-deoxyuridine and 5-methyluridine both displayed low affinity for TgUUT1, with average K_i_ values of 2380 ± 770 µM and 134 ± 31 µM, respectively (Figure 3B). This shows that the 5-methyl group very strongly interferes with the binding of the pyrimidine substrates, and that the 2’-OH group contributes positively to the binding of uridine, likely through a hydrogen bond (δ(ΔG^0^) = 9.0 kJ/mol compared with uridine). The kinetic parameters of TgUUT1 are summarised in Table 1.

**Figure 3.**
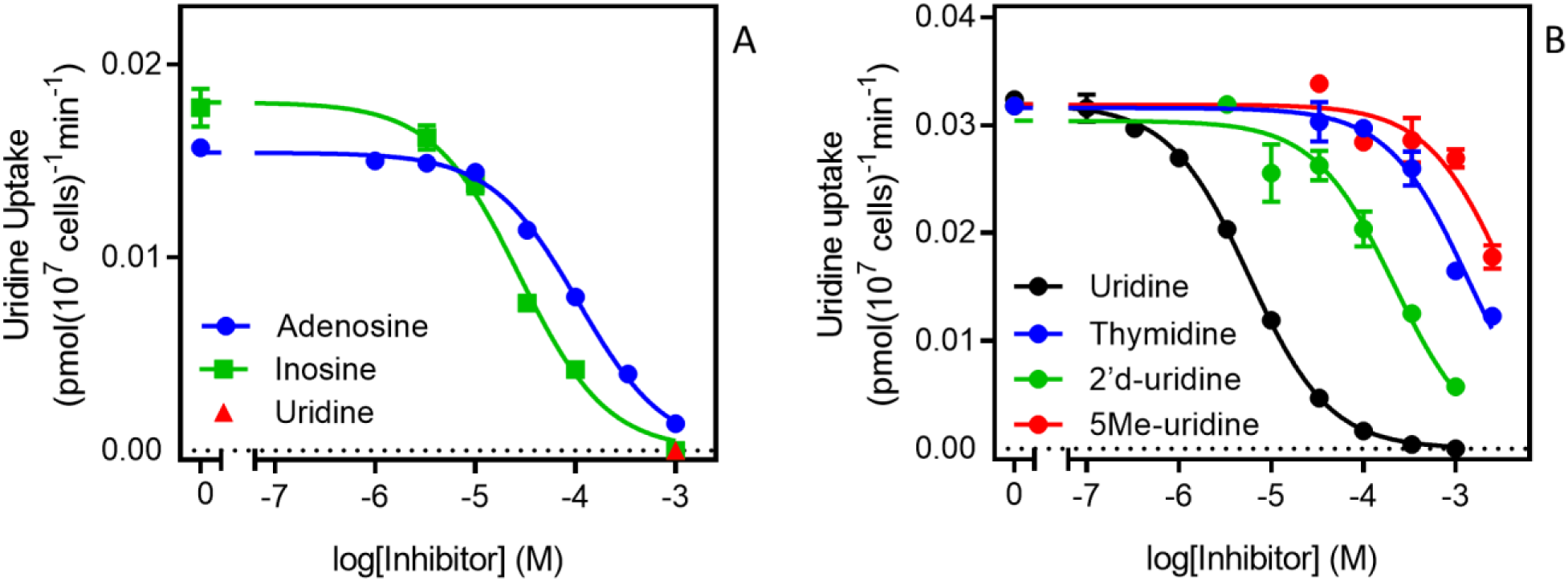
Specificity of the TgUU1 transporter. Inhibition curves for the transport of 0.1 µM [^3^H]-uridine in tachyzoites by (A) purine nucleosides and (B) pyrimidine nucleosides. Graphs show results of a single experiment in triplicate, representative of several similar experiments. Error bars are SEM.

**Table 1.** Kinetic parameters of TgUU1.

|  | Substrate |  |
| --- | --- | --- |
|  | [<sup>3</sup>H]-uridine | [<sup>3</sup>H]-uracil |
| $K_m$<sup>1</sup> | $3.34 \pm 0.82$ | $2.05 \pm 0.40$ |
| $V_{max}$<sup>2</sup> | $1.21 \pm 0.24$ | 0.166 |
| *K<sub>i</sub> values*<sup>1</sup> |  |  |
| Uridine | ---- | $2.44 \pm 0.59$ |
| Uracil | $1.15 \pm 0.07$ | --- |
| 5F-Uracil | $5.24 \pm 0.72$ | $6.80 \pm 2.12$ |
| Thymidine | $1010 \pm 145$ | ND |
| 2'-Deoxyuridine | $134 \pm 31$ | ND |
| 5-Methyluridine | $2380 \pm 774$ | ND |
| Adenosine | $112 \pm 6$ | ND |
| Inosine | $28.2 \pm 4.1$ | ND |

All values are average and Sem of at least three independently conducted experiments, each in triplicate.
<sup>1</sup>, [μM]
<sup>2</sup>, [pmol(10<sup>7</sup> cells)<sup>-1</sup> min<sup>-1</sup>]
ND, not determined

### 3.3 *In vitro* tests of pyrimidine antimetabolites against *T. gondii* tachyzoites in HFF cells

IN the previous section we show that TgUUT1 recognises 5F-pyrimidines as inhibitors, and likely, as substrates. As 5F-pyrimidines are thymidine analogues and are known cancer antimetabolites under the name Floxuridine (Power and Kemeny, 2009; Vodenkova et al., 2020) and have very significant activity against some protozoa (Ali et al., 2013; Alzahrani et al., 2017), we decided to test some of these compounds against intracellular tachyzoites in culture.

Most published protocols for *in vitro* drug screening against *T. gondii* use the MTT assay, which is a colourimetric assay that relies on tetrazolium salt and is used for accessing cell viability, cell proliferation and cytotoxicity for the host cells (Jin et al., 2012). Our protocol is an adaptation from a method that was used to evaluate antiretroviral compounds against *T. gondii* tachyzoites (Wang et al., 2019). These authors added 5×10^4^ cells/mL freshly egressed tachyzoites to each well of a 96-well plate with an HFF cell monolayer, followed by incubation for 4 h to allow the cells to invade the host cells; the medium containing extracellular tachyzoites was then removed and fresh medium containing the desired concentration of the antiretroviral drugs was added to the wells and incubated for 5 days at 37 °C and 5% CO_2_ in a humidified incubator (Wang et al., 2019). In our adaptation of this protocol, we aimed to eliminate the step of wash/removal of tachyzoites, in order to increase reproducibility. We thus prepared the serial dilutions of compounds in the 96-well plates containing the HFF monolayer cells and then added the lower number of 1×10^4^ cells/mL freshly lysed tachyzoites to the 96-well plates, already containing both the HFF monolayer cells and tested compounds. The plates were then incubated for 6 days at 37 °C and 5% CO_2_ in a humidified incubator. A major advantage that was introduced was detection of *T. gondii* in the wells, which was achieved by measuring fluorescence of the RFP-expressing parasites.

We tested our modified protocol using a set of known anti-toxoplasma compounds (sulfadiazine, pyrimethamine, Ara-A and NBMPR) against *T. gondii* in 2 – 4 biological repeats (Figure 4A), and the result was consistent in each repeat. Ara-A has been shown to have moderate activity against *T. gondii* (Pfefferkorn and Pfefferkorn, 1976). Our drug screening assay protocol was consistent with the previous reports as Ara-A showed modest activity against *T. gondii* with EC_50_ 11.4 ± 1.8 µM (n=4). NBMPR has been shown to be very selectively toxic to *T. gondii* with EC_50_ values of 10.2 µM, without apparent cytotoxicity on uninfected HFF monolayer cells up to 100 µM (el Kouni et al., 1999). Our result was similar and reproducible, with EC_50_ of 3.19 ± 0.14 µM (n=4) and no apparent cytotoxicity for NPMPR on uninfected HFF cells at 100 µM. Both purine analogues displayed stronger activity than the current drugs sulfadiazine and pyrimethamine (Figure 4A, Table 2).

**Figure 4.**
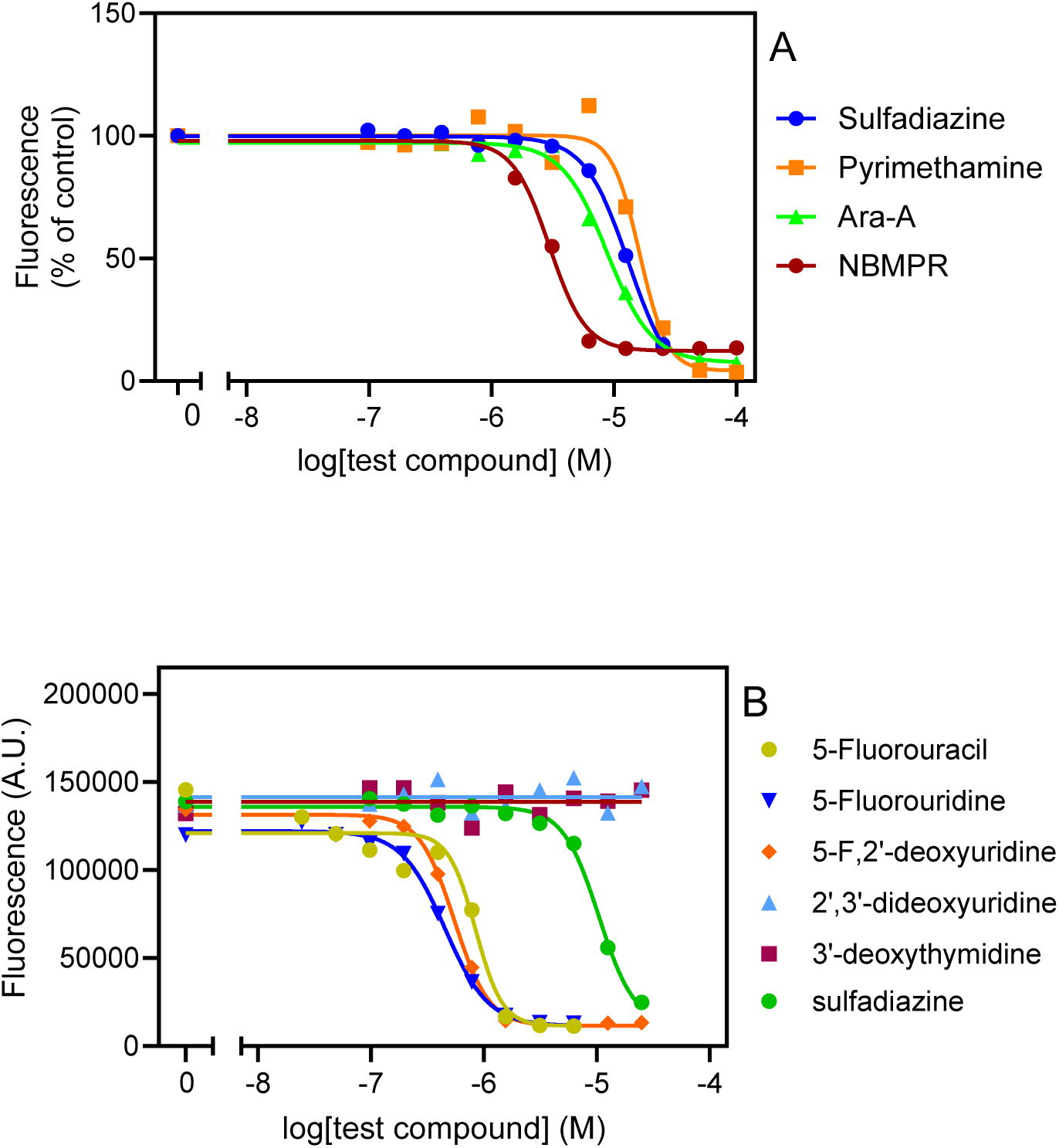
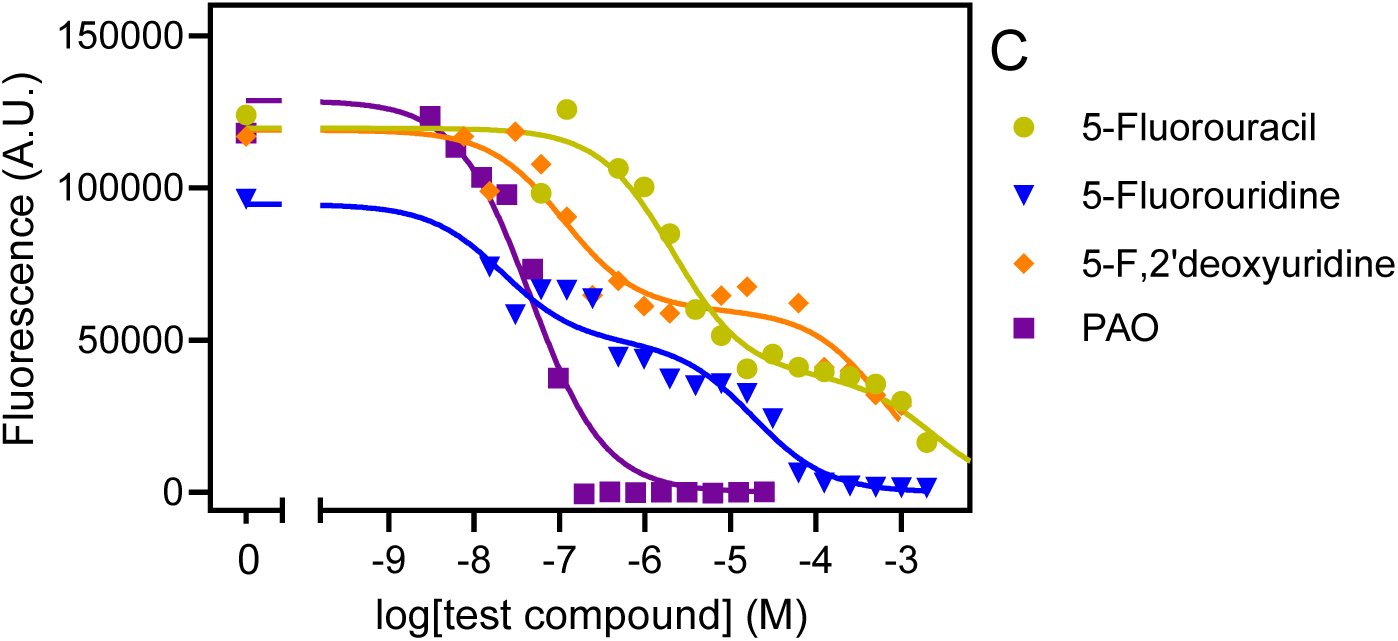
Effect of known anti-toxoplasma agents (A) and pyrimidine analogues (B) on intracellular tachyzoites and on uninfected HFF cells (C). Data were plotted to a log[Inhibitor] vs response sigmoid model with variable slope (A, B) or an equation for two-site competition (C) in Prims 9 to determine EC_50_ values. The curves are single experiments, representative of at least 2 (pyrimethamine) or 3 (all other compounds) independent, identically-performed experiments.

**Table 2.** *In vitro* EC_50_ values of 5-fluoropyrimidines and assorted anti-toxoplasma agents against *T. gondii* intracellular tachyzoites and HFF cells.

| Compound | tachyzoites | HFF ctotoxicity |  |  |
| --- | --- | --- | --- | --- |
|  | EC<sub>50</sub> (μM) | EC<sub>50</sub> (μM)<br>Cytostatic | EC<sub>50</sub> (μM)<br>Cytostatic | S.I. |
| 5-FU | 1.13 ± 0.16 | 2.49 ± 0.43 | 3070 ± 340 | 2720 |
| 5-FUrd | 0.45 ± 0.02 | 0.11 ± 0.10 | 20.4 ± 1 | 45 |
| 5-F,2'dUrd | 0.67 ± 0.07 | 0.19 ± 0.05 | 1480 ± 510 | 2205 |
| Sulfadiazine | 12.4 ± 1.08 | ND | ND | -- |
| Pyrimethamine | 16.2 | ND | ND |  |
| Ara-A | 11.4 ± 1.8 | ND | ND |  |
| NBMPR | 3.19 ± 0.14 | ND | ND |  |
| PAO | ND | ND | 0.06 ± 0.003 | -- |

SI, selectivity index (HFF cytotoxic EC<sub>50</sub>/tachyzoite EC<sub>50</sub>). Each value is the average of at least three independent determinations and the SEM, except for the pyrimethamine EC<sub>50</sub> (n=2). PAO, phenylarsine oxide.

We next tested the 5-F-pyrimidines 5-FU, 5-FUrd and 5-F,2’dUrd for activity against *T. gondii* (Figure 4B). Table 1 shows that the three fluoropyrimidines displayed quite similar EC_50_ values, ranging from 0.45 µM for 5-FUrd to 1.13 µM 5-FU. As such, each of these compounds was at least an order of magnitude more active than sulfadiazine, which was included as a positive control. Other pyrimidine nucleosides tested (3’-deoxythymidine, 2’,3’-dideoxyuridine, 5’-deoxyuridine and 2’-deoxyuridine) had no effect on intracellyular tachyzoites at concentrations up to 25 µM. Next, the three 5-fluoropyrimidines were tested for effects on the uninfected HFF host cells, using the same incubation times and conditions as for the infected cells; Phenylarsine oxide (PAO) was used as the positive control for cytotoxicity. Using this prolonged resazurin-based protocol, the traces for all three 5-fluoropyrimidines were bi-phasic, with low concentration EC_50_ values for growth inhibition (cytostatic) and a second phase yielding (higher) cytocidal EC_50_ values (Figure 4C), much like we previously reported for some nucleoside analogues against *T. brucei* (Rodenko et al., 2007). Using the cytocidal EC_50_s, it was observed that the two clinically used anticancer drugs, 5-FU and 5-F,2’dUrd had a high selectivity (>2000) whereas 5-FUrd was much more toxic and as a result had a poor selecivity index of just 45 (Table 2).

### 3.4 Effects of 5-FU and 5-F2’-dUrd in a mouse model of acute toxoplasmosis

Groups of 6 mice were infected with *T. gondii* by i.p. injection and treated with either 5-FU (25 mg/kg in water, p.i.), 5-F,2’dUrd (25 mg/kg in water, p.i.), sulfadiazine (100 μg/L in water, p.o.) or water (p.i.). The treatments were administered 72 h post infection (p.i.) and thence at 24 h intervals through day 9 p.i., with mice being euthanised with CO_2_ on day 10 p.i. Weight of each mouse was monitored throughout the experiment (Supplemental Figure S1). In the control group, weight decreased slightly over the 10-day period (11.1%, *P* = 0.03, unpaired t-test), and this was less pronounced in each of the treated groups, but none of them had an average weight that was significantly different at the end of the experiment (*P* > 0.1).

The parasite burden was assessed by qPCR, detecting *T. gondii* DNA – specifically the B1 gene (Lin et al., 2000) in lung tissue dissected from the euthanised mice. In all three treated groups, the parasite burden in lung tissue was >85% reduced compared to the untreated control mice (*P* < 0.001; **Figure 5**). 5-F,2’dUrd was significantly less effective, at the chosen dose and administration protocol, than sulfadiazine (*P* < 0.05) but 5-FU treatment was not significantly different from sulfadiazine (*P* > 0.05), but it should be noted that sulfadiazine was administered orally in water, at 40 mg/kg body weight, whereas the two pyrimidines were injected i.p. at 25 mg/kg BW.

**Figure 5.**
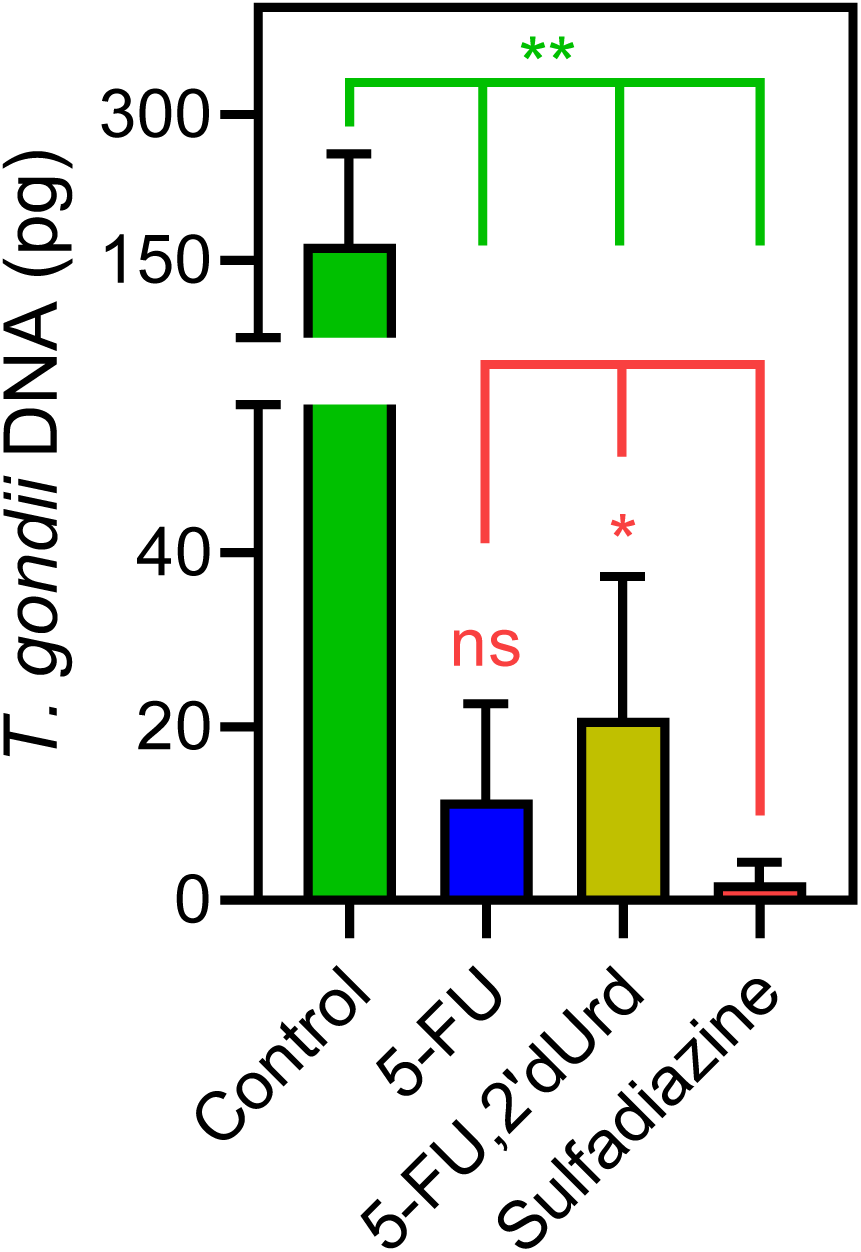
Quantification of *T. gondii* DNA in lung tissue of infected mice, dissected after 10 days of infection with or without treatment, as indicated (n = 6, error bars are SD). *, *P* < 0.05; **, *P* < 0.01 by unpaired, two-tailed t-test.

## 5. DISCUSSION

The first line treatment against acute toxoplasmosis is a combination of pyrimethamine, an inhibitor of dihydrofolate reductase (DHFR) and an inhibitor of dihydropteroate synthetase (DHPS), usually sulfadiazine (Dunay et al., 2018). This combination is only active against acute toxoplasmosis and does not affect cysts in the chronic stage of toxoplasmosis. As pyrimethamine also inhibits human DHFR it is known for its haematological toxicity, making it necessary to give leucovorin or folinic acid together with the first-line treatment to combat the side effect (Alday and Doggett, 2017). Multiple additional disadvantages have been shown for this combination, such as poor tolerance among immunocompromised individuals, toxicity, and a prolonged course of treatment that causes the patients to discontinue taking the medicine, a recent surge in cost, unavailability in some countries, the lack of an intravenous formulation, agranulocytosis, Stevens-Johnson syndrome and hepatic necrosis (Alday and Doggett, 2017; Dunay et al., 2018). Identification of new inhibitors could aid improve treatments in the long term.

The combination of the two dugs pyrimethamine and sulfadiazine disrupts the parasite’s folate synthesis and utilisation (Alday and Doggett, 2017; Dunay et al., 2018). In turn, this results in the inhibition of the nucleotide thymidine monophosphate from 2’-deoxyuridine monophosphate by Thymidylate Synthase, which utilises methylene-tetrahydrofolate as its methyl donor. *T. gondii* is particularly vulnerable to drugs targeting this pathway because it, unusually, lacks a thymidine kinase activity (Fox et al., 2001) and is therefore unable to incorporate any salvaged thymidine into its nucleotide pool. However, *T. gondii* is known to be able to grow on extracellular uridine or uracil as its only pyrimidine source (Fox and Bzik, 2002; Fox and Bzik, 2010). It is further likely that cytidine can be salvaged by *T. gondii* and converted to uridine by cytidine deaminase (Iltzsch, 1993), if the parasite can acquire extracellular cytidine. These observations led us to hypothesise that *T. gondii* tachyzoites may have one or more carriers for uracil and uridine, and possibly for cytidine, but may not possess capacity to take up thymidine.

We tested the transport by tachyzoites of uracil, uridine, cytidine and thymidine. Only the accumulation of 0.25 µM [^3^H]-thymidine was, though just-measurable over 10 minutes, was not saturable, in that it was not inhibited by a 4000-fold excess (1 mM) of unlabelled thymidine, or by 250 µM uridine. This clearly establishes that *T. gondii* tachyzoites do not express a thymidine-specific thymidine transporter, although it cannot be ruled out that they express a transporter for the uptake of a different substrate, that could also have some capacity to internalise thymidine, with very low affinity and efficiency. [^3^H]-Cytidine uptake was also quite slow but 100% inhibited by 1 mM unlabelled cytidine and by 250 µM uridine, revealing the expression of cytidine transporter that is highly sensitive to inhibition by uridine. Uptake of [^3^H]-uridine was relatively robust and displayed similarly high affinity for uridine and for uracil as an inhibitor. The same was observed using [^3^H]-uracil as substrate and uridine as inhibitor, establishing reciprocal inhibition with statistically identical K_m_ and K_i_ values. Moreover, uridine and uracil both inhibited [^3^H]-uridine and [^3^H]-uracil completely and with Hill slopes that were consistently very close to −1. These are all parameters indicating a single transporter for uridine and uracil and their really is no reasonable alternative explanation for all the data; we have designated this *T. gondii* pyrimidine carrier uridine/uracil transporter 1 (TgUUT1). It is likely that this carrier also has a minor capacity to take up cytidine, but this is likely to be of less relevance physiologically due to the very poor rate of uptake, the low availability of intracellular free cytidine and the fact that cytidine needs to be degraded to uridine, and subsequently to uracil, by the parasite.

A high affinity uridine/uracil transporter was recently also described in *Leishmania mexicana* (LmexUUT1), but that transporter also had high affinity for adenosine (Alzahrani et al., 2017). This is not the case, however, for TgUUT1, as it was inhibited only ∼70% by a 2500-fold excess of unlabelled adenosine (Figure 1C) and the adenosine K_i_ for uridine transport was determined at 112 µM. Nor is TgUUT1 similar to the uracil transporters of other protozoa, as the uracil transporters of *T. brucei* (De Koning and Jarvis, 1998; Gudin et al., 2006), *L. major* (Papageorgiou et al., 2005) and *T. vaginalis* (Natto et al., 2021b) are all highly specific for uracil and exclude uridine, which is taken up by separate transporters by those species (Aldfer et al., 2022; Ali et al., 2013; Natto et al., 2021b).

The non-specificity of TgUUT1 regarding uracil and uridine would suggest that the ribose moiety of uridine is not engaged in interactions with the transporter but essentially a tolerated substitution on N1 of the pyrimidine ring – much like the TbAT1/P2 transporter of *T. brucei*, which has only a marginally higher affinity for adenine than adenosine although adenine is translocated with higher efficiency as seen from the V_max_/K_m_ (1.84 *vs* 1.32; De Koning and Jarvis, 1999), making TbAT1 essentially an adenine transporter that tolerates adenosine. Recently, we reported that the *T. gondii* transporter Tg_244441 also transported both nucleobases and nucleosides with similar affinity, albeit in this case oxopurines rather than the aminopurines taken up by TbAT1/P2 (Campagnaro et al., 2022). However, in this case, the ribose moiety was not carried passively through the transporter; instead the similar affinity was in this case the result of a different set of interactions for nucleobases and nucleosides adding up to similar levels of binding energy.

For the TgUU1 transporter reported here, the so far limited evidence would suggest that the ribose moiety, and specifically the 2’-OH group, does contribute to the binding of uridine: 2’-deoxyuridine displayed a K_i_ value of 134 ± 31 µM, compared to the uridine K_m_ of 3.34 ± 0.82 µM (*P* < 0.01) – a loss of 9.1 kJ/mol in Gibbs free energy of interaction (δ(ΔG^0^) = 9.1 kJ/mol). This observation goes some way to explain the very low affinity for thymidine, which also lacks 2’-OH and displays a K_i_ of 1010 ± 145 µM, corresponding to δ(ΔG^0^) of 14.2 kJ/mol. The greater loss of affinity for thymidine than for 2’-dUrd must be attributed to the 5-methyl group, which is the only difference between the molecules. Consistent with this interpretation 5-methyluridine displayed very low affinity and it can be concluded that the 5-methyl substitution is not compatible with pyrimidine nucleoside transport by TgUUT1. The much smaller 5-fluoro substitution, however, was much better tolerated as the δ(ΔG^0^) amounted to only 3.0 kJ/mol (5-FU K_i_ compared to uracil K_m_), allowing the uptake of 5F-pyrimidines.

5-F-pyrimidines have been mentioned as potential therapeutic agents before. As early as 1977, Pfefferkorn and Pfefferkorn (1977b) reported that 5-F,2’dUrd inhibited *T. gondii* growth in vitro and rapidly inhibited its nucleic acid synthesis. A 5-F,2’dUrd resistant cell line was created and found to lack UPRT activity (Pfefferkorn, 1978). Subsequently, it was found that 5-F-pyrimidines are good substrates for uridine phosphorylase (Iltzsch and Klenk, 1993). This observation, and our finding of equal transport of uridine and uracil, together explains why we found virtually equal activity of 5-FUrd, 5-F,2’dUrd and 5-FU, as the nucleosides are both converted to 5-FU by uridine phosphorylase inside the parasite, prior to activation to 5-FUMP by UPRT.

The *in vitro* activity of the 5-F-pyrimidines being between 11 and 27-fold stronger than first-line drug sulfadiazine, and no data on toxoplasmosis in an animal model having been reported for these compounds, we conducted a first trial in a standardised mouse model with 5-FU and 5-F,2’dUrd; the S.I. of 5FUrd was deemed too low. The parasite load was quantified with qPCR and compared to groups of mice treated with sulfadiazine or vehicle. The length of this initial infection was ten days, limiting the test to acute toxoplasmosis (Lin et al., 2000). While the differences in dosage regime between sulfadiazine (p.o., 100 mg/kg ×7) and the 5-F-pyrimidines (i.p., 25 mg/kg ×7) makes a genuine quantitative comparison of relative efficacy tenuous, all three treatments were similarly effective in reducing the parasite burden compared to the vehicle control. This invites further investigations into 5-FU, particularly, as an anti-toxoplasmosis agent, especially since it is already in long-time clinical use as an anti-cancer drug (Vodenkova et al., 2020). Interestingly, 5-FU has already been used in one study to treat AIDS patients with cerebral toxoplasmosis, in combination with clindamycin (Dhiver et al., 1993), a second-line treatment for toxoplasmosis (Konstantinovic et al., 2019). The brief report concluded that 5-FU/clindamycin is and effective and less toxic alternative to pyrimethamine/sulfadiazine, the toxicity of 5-FU being limited because ‘an effective anti-toxoplasma activity is obtained with doses tenfold less than those used for cancer chemotherapy.’

In summary, we studied the uptake of pyrimidine nucleosides and uracil in *T. gondii* and report here a novel uridine/uracil carrier, TgUUT1, which also mediates the uptake of 5-fluoropyrimidine antimetabolites, but not of thymidine. This finding strongly contributes to our understanding of pyrimidine metabolism in *T. gondii* and explains the basis for the strong anti-toxoplasma activity of 5-F-pyrimidines. Further *in vivo* studies with 5-FU, in particular, seem warranted, as well as further work on the pharmacologically relevant *T. gondii* nucleoside transporters.

## Acknowledgements

H.A.A.E. was supported by a studentship from the government of Libya. We thank Mr Andrew Boyd for expert assistance with the cell cultures.

## References

Al-Salabi, M.I., and De Koning, H.P. (2005). ‘Purine nucleobase transport in amastigotes of *Leishmania mexicana*: involvement in allopurinol uptake.’. Antimicrob. Agents Chemother. 49, 3682–3689.

Al Safarjalani, O.N., Rais, R.H., Kim, Y.A., Chu, C.K., Naguib, F.N., and El Kouni, M.H. (2010). ‘Carbocyclic 6-benzylthioinosine analogues as subversive substrates of *Toxoplasma gondii* adenosine kinase: Biological activities and selective toxicities.’. Biochem. Pharmacol. 80, 955– 963.

Alday, P. H., and Doggett, J. S. (2017). ‘Drugs in development for toxoplasmosis: advances, challenges, and current status.’. Drug Des. Dev. Ther. 11, 273.

Aldfer, M., Alfayez, I.A., Elati, H.A.A., Gayen, N., Elmahallawy, E.K., Murillo, A.M., Marsiccobetre, S., Van Calenbergh, S., Silber, A.M., and De Koning, H.P. (2022). ‘The *Trypanosoma cruzi* nucleoside transporter TcrNT2 is a conduit for the uptake of 5-F-2’deoxyuridine and tubercidin analogues.’. Molecules 27, 8045.

Aldfer, M.M., AlSiari, T.A., Elati, H.A.A., Natto, M.J., Alfayez, I.A., Campagnaro, G.D., Sani, B., Burchmore, R.J.S., Diallinas, G., and De Koning, H.P. (2022). ‘Nucleoside transport and nucleobase uptake null mutants in *Leishmania mexicana* for the routine expression and characterisation of purine and pyrimidine transporters.’. Int. J. Mol. Sci. 23, 8139.

Ali, J.A.M., Creek, D.J., Burgess, K., Allison, H.C., Field, M.C., Mäser, P., and De Koning, H.P. (2013). ‘Pyrimidine salvage in *Trypanosoma brucei* bloodstream forms and the trypanocidal action of halogenated pyrimidines.’. Mol. Pharmacol. 83, 439–453.

Alzahrani, K.J.H., Ali, J.A.M., Eze, A.A., Looi, W.L., Tagoe, D.N.A., Creek, D.J., Barrett, M.P., and De Koning, H.P. (2017). ‘Functional and genetic evidence that nucleoside transport is highly conserved in *Leishmania* species: implications for pyrimidine-based chemotherapy.’ Int. J. Parasitol. Drugs Drug Resist. 7, 206–226.

Berens, R. L., Krug, E. C., and Marr, J. J. (1995). ‘Purine and pyrimidine metabolism.’. In: Biochemistry and molecular biology of parasites. Joseph J. Marr, and Miklos Muller, eds. Elsevier. pp. 89–118.

Campagnaro, G. D., and De Koning, H. P. (2020). ‘Purine and pyrimidine transporters of pathogenic protozoa – conduits for therapeutic agents.’. Med. Res. Rev. 40, 1679–1714.

Campagnaro, G.D., Elati, H.A.A., Balaska, S., Martin Abril, M.E., Natto, M.J., Hulpia, F., Lee, K., Sheiner, L., Van Calenbergh, S., and De Koning, H.P. (2022). ‘A *Toxoplasma gondii* oxopurine transporter binds nucleobases and nucleosides using different binding modes.’. Int. J. Mol. Sci. 23, 710.

Cheviet, T., Lefebvre-Tournier, I., Wein, S., and Peyrottes, S. (2019). ‘*Plasmodium* purine metabolism and its inhibition by nucleoside and nucleotide analogues.’. J. Med. Chem. 62, 8365–8391.

Chiang, C.W., Carter, N., Sullivan, W.J. Jr., Donald, R.G., Roos, D.S., Naguib, F.N., el Kouni, M.H., Ullman, B., and Wilson, C.M. (1999). ‘The adenosine transporter of *Toxoplasma gondii*. Identification by insertional mutagenesis, cloning, and recombinant expression.’. J. Biol. Chem.274, 35255–35261.

De Koning, H. P., Al-Salabi, M. I., Cohen, A. M., Coombs, G. H., and Wastling, J. M. (2003). ‘Identification and characterisation of high affinity nucleoside and nucleobase transporters in *Toxoplasma gondii*.’. Int. J. Parasitol. 33, 821–831.

De Koning, H. P., and Jarvis, S. M. (1998). ‘A highly selective, high-affinity transporter for uracil in *Trypanosoma brucei brucei*: evidence for proton-dependent transport.’. Biochem. Cell Biol. 76, 853–858.

De Koning, H.P., and Jarvis, S.M. (1999) ‘Adenosine transporters in bloodstream forms of *T. b. brucei*: Substrate recognition motifs and affinity for trypanocidal drugs.’. Mol. Pharmacol. 56, 1162–1170.

Deniskin, R., Frame, I.J., Sosa, Y., and Akabas, M.H. (2015). ‘Targeting the *Plasmodium vivax* equilibrative nucleoside transporter 1 (PvENT1) for antimalarial drug development.’. Int. J. Parasitol. Drugs Drug Resist. 6, 1–11.

Dhiver, C., Milandre, C., Poizot-Martin, I., Drogoul, M. P., Gastaut, J. L., and Gastuat, J. A. (1993). ‘5-Fluoro-uracil-clindamycin for treatment of cerebral toxoplasmosis.’. Aids 7, 143–144.

Dunay, I.R., Gajurel, K., Dhakal, R., Liesenfeld, O., and Montoya, J.G. (2018). ‘Treatment of toxoplasmosis: historical perspective, animal models, and current clinical practice.’. Clin. Microbiol. Rev. 31, e00057–17.

Finley, R. W., Cooney, D. A., and Dvorak, J. A. (1988). ‘Nucleoside uptake in *Trypanosoma cruzi*: analysis of a mutant resistant to tubercidin.’. Mol. Biochem. Parasitol. 31, 133–140.

Fiuza, L.F.A., Batista, D.G., Girão, R.D., Hulpia, F., Finamore-Araújo, P., Aldfer, M.M., Elmahallawy, E.K., De Koning, H.P., Moreira, O., Van Calenbergh, S., and Soeiro, M.D. (2022). ‘Phenotypic evaluation of nucleoside analogues against *Trypanosoma cruzi* infection: *In vitro* and *in vivo* approaches.’. Molecules 27, 8087.

Fox, B. A., Belperron, A. A., and Bzik, D. J. (2001). ‘Negative selection of herpes simplex virus thymidine kinase in *Toxoplasma gondii*.’ Mol. Biochem. Parasitol. 116, 85–88.

Fox, B. A., and Bzik, D. J. (2002). ‘De novo pyrimidine biosynthesis is required for virulence of *Toxoplasma gondii*.’. Nature 415, 926–929.

Fox, B. A., and Bzik, D. J. (2010). ‘Avirulent uracil auxotrophs based on disruption of orotidine-5′-monophosphate decarboxylase elicit protective immunity to *Toxoplasma gondii*.’. Infect. Immun. 78, 3744–3752.

Fox, B. A., and Bzik, D. J. (2020). ‘Biochemistry and metabolism of *Toxoplasma gondii*: purine and pyrimidine acquisition in Toxoplasma gondii and other Apicomplexa.’. In: Weiss, L. M. and Kim, K. (eds.) Toxoplasma gondii. Third edition ed. San Diego: Elsevier Academic Press.

Ghosh, M., and Mukherjee, T. (2000). ‘Stage-specific development of a novel adenosine transporter in *Leishmania donovani* amastigotes.’. Mol. Biochem. Parasitol. 108, 93–99.

Gudin, S., Quashie, N.B., Candlish, D., Al-Salabi, M.I., Jarvis, S.M., Ranford-Cartwright, L.C., and De Koning, H.P. (2006). ‘*Trypanosoma brucei*: A survey of pyrimidine transport activities.’. Exp. Parasitol. 114, 103–108.

Gutteridge, W.E., and Trigg, P.I. (1970). ‘Incorporation of radioactive precursors into DNA and RNA of *Plasmodium knowlesi in vitro*.’. J. Protozool. 17, 89–96.

Hassan, H. F., and Coombs, G. H. (1988). ‘Purine and pyrimidine metabolism in parasitic protozoa.’. FEMS Microbiol. Lett. 54, 47–83.

Heppler, L. N., Attarha, S., Persaud, R., Brown, J. I., Wang, P., Petrova, B., et al. (2022). ‘The antimicrobial drug pyrimethamine inhibits STAT3 transcriptional activity by targeting the enzyme dihydrofolate reductase.’. J. Biol. Chem. 298, 101531.

Hill, D. E., Chirukandoth, S., and Dubey, J. P. (2005). ‘Biology and epidemiology of *Toxoplasma gondii* in man and animals.’. Anim. Health Res. Rev. 6, 41–61.

Hulpia, F., Bouton, J., Campagnaro, G.D., Alfayez, I.A., Mabille, D., Maes, L., De Koning, H.P., Caljon, G., and Van Calenbergh, S. (2020a). ‘C6-O-Alkylated 7-deazainosine nucleoside analogues: Discovery of potent and selective anti-sleeping sickness agents.’. Eur. J. Med. Chem. 188, 112018.

Hulpia, F., Campagnaro, G.D., Alzahrani, K.J., Alfayez, I.A., Ungogo, M.A., Mabille, D., Maes, L., De Koning, H.P., Caljon, G., and Van Calenbergh, S. (2020). ‘Structure-activity relationship exploration of 3’-deoxy-7-deazapurine nucleoside analogues as anti-*Trypanosoma brucei* agents.’. ACS Infect. Dis. 6, 2045–2056.

Iltzsch, M. H. (1993). ‘Pyrimidine salvage pathways in *Toxoplasma gondii*.’. J. Euk. Microbiol. 40, 24– 28.

Iltzsch, M. H., and Klenk, E. E. (1993). ‘Structure-activity relationship of nucleobase ligands of uridine phosphorylase from *Toxoplasma gondii*.’. Biochem. Pharmacol. 46, 1849–1858.

Jin, C., Jung, S.Y., Kim, S.Y., Song, H.O., and Park, H. (2012). ‘Simple and efficient model systems of screening anti-*Toxoplasma* drugs *in vitro*.’. Expert Opin. Drug Disc. 7, 195–205.

Kim, Y.A., Rawal, R.K., Yoo, J., Sharon, A., Jha, A.K., Chu, C.K., Rais, R.H., Al Safarjalani, O.N., Naguib, F.N., and El Kouni, M.H. (2010). ‘Structure–activity relationships of carbocyclic 6-benzylthioinosine analogues as subversive substrates of *Toxoplasma gondii* adenosine kinase.’. Bioorg. Med. Chem. 18, 3403–3412.

Konstinovic, N., Guegan, H., Stajnär, T., Belaz, S., and Robert-Gangneux, F. (2019). ‘Treatment of toxoplasmosis: Current options and future perspectives.’. Food Waterborne Parasitol. 15, e00036.

Lin, M.H., Chen, T.C., Kuo, T.T., Tseng, C.C., and Tseng, C.P. (2000). ‘Real-time PCR for quantitative detection of *Toxoplasma gondii*.’. J. Clin. Microbiol. 38, 4121–4125.

Mabille, D., Ilbeigi, K., Hendrickx, S., Ungogo, M.A., Hulpia, F., Lin, C., Maes, L., De Koning, H.P., Van Calenbergh, S., and Caljon, G. (2022). ‘Nucleoside analogues for the treatment of animal trypanosomiasis.’. Int. J. Parasitol. Drugs Drug Resist. 19, 21–30.

Montoya, J., and Liesenfeld, O. (2004). ‘Toxoplasmosis.’. Lancet 363, 1965–1976.

Natto, M.J., Hulpia, F., Kalkman, E.R., Baillie, S., Alhejeli, A., Miyamoto, Y., Eckmann, L., Van Calenbergh, S., and De Koning, H.P. (2021a). ‘Deazapurine nucleoside analogues for the treatment of *Trichomonas vaginalis*.’. ACS Infect. Dis. 7, 1752–1764.

Natto, M.J, Miyamoto, Y., Munday, J.C., AlSiari, T.A., Al-Salabi, M.I., Quashie, N.B., Eze, A.A., Eckmann, L., and De Koning, H.P. (2021b). ‘Comprehensive characterization of purine and pyrimidine transport activities in *Trichomonas vaginalis* and functional cloning of a trichomonad nucleoside transporter.’. Mol. Microbiol. 116, 1489–1511.

Papageorgiou, I.G., Yakob, L., Al-Salabi, M.I., Diallinas, G., Soteriadou, K., and De Koning, H.P. (2005). ‘Identification of the first pyrimidine nucleobase transporter in *Leishmania*: similarities with the *Trypanosoma brucei* U1 transporter and antileishmanial activity of uracil analogues.’. Parasitology 130, 275–283.

Parker, M.D., Hyde, R.J., Yao, S.Y., McRobert, L., Cass, C.E., Young, J.D., McConkey, G.A., and Baldwin, S.A. (2000). ‘Identification of a nucleoside/nucleobase transporter from *Plasmodium falciparum*, a novel target for anti-malarial chemotherapy.’. Biochem. J. 349, 67–75.

Polet, H., and Barr, C.F. (1968). ‘DNA, RNA, and protein synthesis in erythrocytic forms of *Plasmodium knowlesi*.’. Am. J. Trop. Med. Hyg. 17, 672–679.

Power, D.G., and Kemeny, N.E. (2009). ‘The role of floxuridine in metastatic liver disease.’. Mol. Cancer Ther. 8, 1015–1025.

Pfefferkorn, E., and Pfefferkorn, L. C. (1976). ‘Arabinosyl nucleosides inhibit *Toxoplasma gondii* and allow the selection of resistant mutants.’. J. Parasitol. 62, 993–999.

Pfefferkorn, E.R., and Pfefferkorn, L.C. (1977a). ‘Specific labelling of intracellular *Toxoplasma gondii* with uracil.’. J. Protozool. 24, 449–453.

Pfefferkorn, E.R., and Pfefferkorn, L.C. (1977b). ‘*Toxoplasma gondii*: characterisation of a mutant resistant to 5-fluorodeoxyuridine.’. Exp. Parasitol. 42, 44–55.

Pfefferkorn, E.R. (1978). ‘*Toxoplasma gondii*: the enzymic defect of a mutant resistant to 5-fluorodeoxyuridine.’ Exp. Parasitol. 44, 26–35.

Pfefferkorn, E. (1988). ‘The biology of parasitism. A molecular and immunological approach.’. In: Toxoplasma gondii viewed from a virological perspective (Englund, P. T.; Sher, A., Editors). Alan R. Liss, Inc., New York, USA. pp 479–501.

Quashie, N. B., Dorin-Semblat, D., Bray, P. G., Biagini, G. A., Doerig, C., Ranford-Cartwright, L. C., and De Koning, H. P. (2008). ‘A comprehensive model of purine uptake by the malaria parasite *Plasmodium falciparum*: identification of four purine transport activities in intraerythrocytic parasites.’. Biochem. J. 411, 287–295.

Rider Jr, S. D., and Zhu, G. (2010). ‘*Cryptosporidium*: genomic and biochemical features.’ Exp. Parasitol. 124, 2–9.

Rodenko, B., Van der Burg, A.M., Wanner, M.J., Kaiser, M., Brun, R., Gould, M.K., De Koning, H.P., and Koomen, G.J. (2007). ‘2,N6-Disubstituted adenosine analogues with antitrypanosomal and antimalarial activity. Synthesis, uptake studies and *in vivo* evaluation.’. Antimicrob. Agents Chemother. 51, 3796–3802.

Saeij, J. P., Boyle, J. P., Grigg, M. E., Arrizabalaga, G., and Boothroyd, J. C. (2005). ‘Bioluminescence imaging of *Toxoplasma gondii* infection in living mice reveals dramatic differences between strains.’. Infect. Immun. 73, 695–702.

Schwab, J. C., Afifi, M. A., Pizzorno, G., Handschumacher, R. E., and Joiner, K. A. (1995). ‘*Toxoplasma gondii* tachyzoites possess an unusual plasma membrane adenosine transporter.’. Mol. Biochem. Parasitol. 70, 59–69.

Sheiner, L., Demerly, J.L., Poulsen, N., Beatty, W.L., Lucas, O., Behnke, M.S., White, M.W., and Striepen, B. (2011). ‘A systematic screen to discover and analyze apicoplast proteins identifies a conserved and essential protein import factor.’. PLoS Pathog. 7, e1002392.

Tenter, A. M., Heckeroth, A. R., and Weiss, L. M. (2000). ‘*Toxoplasma gondii*: from animals to humans.’. Int. J. Parasitol. 30, 1217–1258.

Ungogo, M.A., Aldfer, M.M., Natto, M.J., Zhang, H., Chisholm, R., Walsh, K., McGee, M.C., Ilbeigi, K., Asseri, J.I., Burchmore, R.J.S., Caljon, G., Van Calenbergh, S., and De Koning, H.P. (2023). ‘Cloning and characterisation of *Trypanosoma congolense* and *T. vivax* nucleoside transporters reveal the potential of P1-type carriers for the discovery of broad-spectrum nucleoside-based therapeutics against Animal African Trypanosomiasis.’. Int. J. Mol. Sci. 24, 3144.

Vizcarra, E.A., Goerner, A.L., Ulu, A., Hong, D.D., Bergersen, K.V., Talavera, M.A., Le Roch, K., Wilson, E.H., and White, M.W. (2023). ‘An *ex vivo* model of *Toxoplasma* recrudescence reveals developmental plasticity of the bradyzoite stage.’. *mBio*, e01836–23.

Vodenkova, S., Buchler, T., Cervena, K., Veskrnova, V., Vodicka, P., Vymetalkova, V. (2020). ‘5-fluorouracil and other fluoropyrimidines in colorectal cancer: Past, present and future.’. Pharmacol. Ther. 206, 107447.

Wang, J.L., Elsheikha, H.M., Li, T.T., He, J.J., Bai, M.J., Liang, Q.L., Zhu, X.Q., and Cong, W. (2019). ‘Efficacy of antiretroviral compounds against *Toxoplasma gondii* in vitro.’. Int. J. Antimicrob. Agents 54, 814–819.

