## Supplementary figures and images for "Pyrimidine salvage in *Toxoplasma gondii* as a target for new treatment"

### Supplemental Figure S1

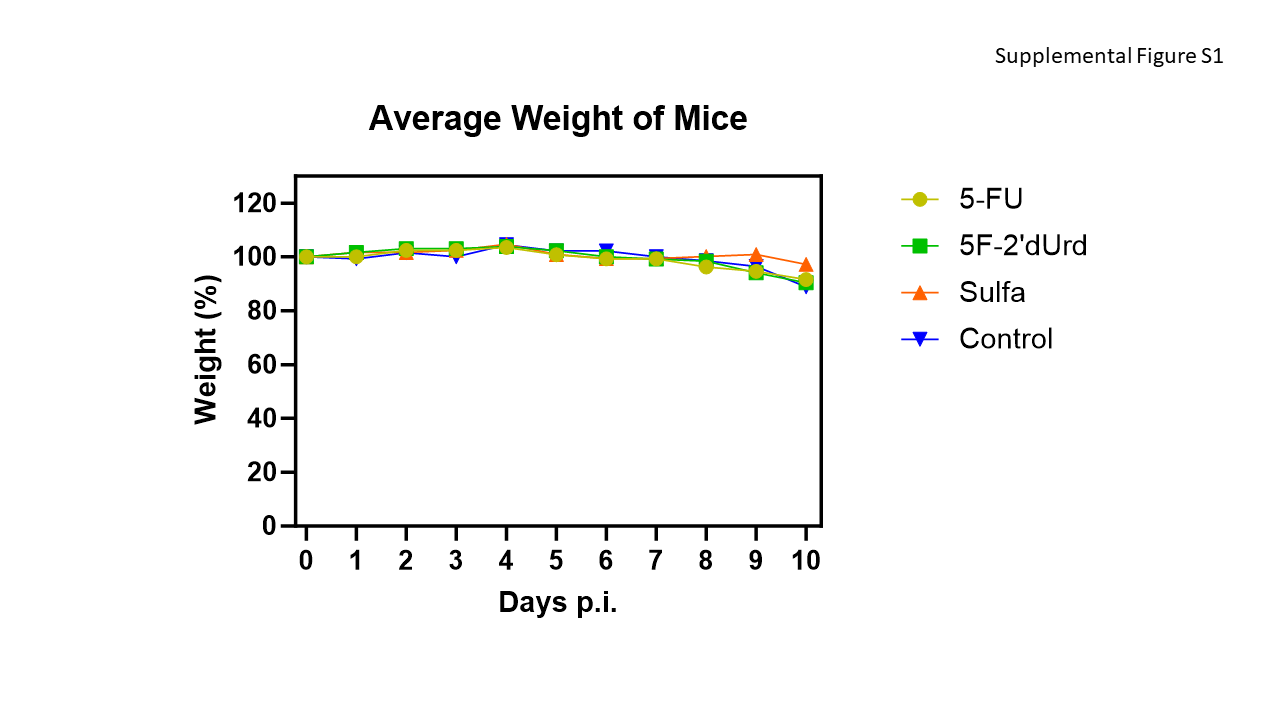
